# Geometric Potentials from Deep Learning Improve Prediction of CDR H3 Loop Structures

**DOI:** 10.1101/2020.02.09.940254

**Authors:** Jeffrey A. Ruffolo, Carlos Guerra, Sai Pooja Mahajan, Jeremias Sulam, Jeffrey J. Gray

## Abstract

Antibody structure is largely conserved, except for a complementarity-determining region featuring six variable loops. Five of these loops adopt canonical folds which can typically be predicted with existing methods, while the remaining loop (CDR H3) remains a challenge due to its highly diverse set of observed conformations. In recent years, deep neural networks have proven to be effective at capturing the complex patterns of protein structure. This work proposes DeepH3, a deep residual neural network that learns to predict inter-residue distances and orientations from antibody heavy and light chain sequence. The output of DeepH3 is a set of probability distributions over distances and orientation angles between pairs of residues. These distributions are converted to geometric potentials and used to discriminate between decoy structures produced by RosettaAntibody. When evaluated on the Rosetta Antibody Benchmark dataset of 49 targets, DeepH3-predicted potentials identified better, same, and worse structures (measured by root-mean-squared distance [RMSD] from the experimental CDR H3 loop structure) than the standard Rosetta energy function for 30, 13, and 6 targets, respectively, and improved the average RMSD of predictions by 21.3% (0.48 Å). Analysis of individual geometric potentials revealed that inter-residue orientations were more effective than inter-residue distances for discriminating near-native CDR H3 loop structures.

## Introduction

The adaptive immune system of vertebrates is responsible for coordinating highly specific responses to pathogens. In such a response, B cells of the adaptive immune system secrete antibodies to bind and neutralize some antigen. The central role of antibodies in adaptive immunity makes them attractive for the development of new therapeutics. However, rational design of antibodies is hindered by the difficulty of experimental determination of macromolecular structures in a high-throughput manner. Advances in computational modeling of antibody structures provides an alternative to experiments, but computations are not yet sufficiently accurate and reliable.

Antibody structure consists of two sets of heavy and light chains that form a highly conserved framework region (F_c_) and two variable regions responsible for antigen binding (F_v_). The structural conservation of the Fc is functionally significant, enabling the recognition of different antibody isotypes by their receptors, and the Fc lends well to homology modeling. The F_v_ contains several segments of sequence hypervariability that provide the structural diversity necessary to bind a variety of antigens. This diversity is largely focused in six β-strand loops known as the complementarity determining regions (CDRs). Five of these loops (L1-L3, H1, H2) typically fold into one of several canonical conformations [1] that are predicted well by existing methods [2]. However, the third CDR loop of the heavy chain (H3) is observed in a diverse set of conformations and remains a challenge to model [3-9].

Application of deep learning techniques has yielded significant advances in the prediction of protein structure in recent years. At CASP13, AlphaFold [10] and RaptorX [11] demonstrated that inter-residue distances could be accurately learned from sequence and coevolutionary features. Both approaches used deep residual network architectures with dilated convolutions to predict inter-residue distances, which provide a more complete structural description than contacts alone. trRosetta built on this progress by expanding beyond distances to predict a set of inter-residue orientations [12]. This rich set of inter-residue geometries allows trRosetta to outperform leading approaches on the CASP13 dataset, even with a shallower network [12].

This work expands on the progress in general protein structure prediction by applying similar techniques to a challenging problem in antibody structure prediction. Specifically, we propose DeepH3, a deep residual network that learns to predict inter-residue distances and orientations from antibody heavy and light chain sequence alone. We show that when compared to the Rosetta energy function, DeepH3-predicted geometric potentials can more accurately identify near-native CDR H3 loop structures.

## Methods

### Overview

DeepH3 is a deep residual network that learns to predict inter-residue distances and orientations from antibody heavy and light chain sequences. The architecture of DeepH3 draws inspiration from RaptorX [11, 13], which performed well on general protein structure prediction at CASP13. The outputs of DeepH3 are converted into geometric potentials in order to better discriminate between CDR H3 loop structure decoys generated using a standard homology modeling approach [14].

### Antibody Structure Datasets

#### Benchmark Dataset

The Rosetta antibody benchmark dataset consists of 49 F_v_ structures with CDR H3 loop lengths ranging from 9 to 20 residues [14, 15]. These structures were selected from the PyIgClassify database [16] based on their quality, with each having resolution of 2.5 Å or better, a maximum *R* value of 0.2, and a maximum *B* factor of 80.0 Å^2^ for every atom [14, 15]. The diversity of the set is enhanced by ensuring that no two structures share a common CDR H3 loop sequence, but limited by restriction to structures from humans and mice [14, 15].

#### Training Dataset

The training dataset for this work was extracted from SAbDab, a curated database of all antibody structures in the Protein Data Bank [17]. We enforced thresholds of 99% sequence identity and 3.0 Å resolution to produce a balanced, high-quality dataset. This high sequence identity cutoff was chosen due to the high conservation of sequence characteristic of antibodies. In cases where multiple chains existed for the same structure, only the first chain in the PDB file was used. Finally, any structures present in the Rosetta antibody benchmark dataset were removed. These steps resulted in 1,462 structures, of which a random 95% were used for model training and 5% were used for validation. This small validation set was found to be sufficient to control for overfitting. Note that testing is carried out on an independent benchmark sharing no structures with the training/validation sets.

### Learning Inter-Residue Geometries from Antibody Sequence

#### Input Features

Unlike most comparable networks, DeepH3 relies only on amino acid sequence as input. For general protein structure prediction, current methods typically utilize some combination of multiple sequence alignments, sequence profiles, co-evolutionary data, secondary structures, *etc.* [10-13, 18]. While these additional input features provide rich information for general protein structure predictions, the highly conserved antibody structure should render these features less useful, and we omit them. DeepH3 takes as input a one-hot encoded sequence formed by concatenating the target heavy and light chains. A chain delimiter is added to the last position in the heavy chain, resulting in an input of dimension Lx21, where L is the cumulative length of the heavy and light chain sequences.

#### Inter-Residue Geometries

In addition to inter-residue distances, DeepH3 is also trained to predict the set of dihedral and planar angles previously proposed for trRosetta [12]. For two residues *i* and *j*, the relative orientation is defined by six parameters (*d, ω*, θ_*ij*_, θ_*ji*_, *φ*_*ij*_, and *φ*_ji_, Figure 1A-B, adapted from [12]). The distance (d) is defined using C_β_ atoms or for glycine residues, C_α_. Distances were discretized into 26 bins, with 24 in the range of [4, 16 Å] and two additional bins for all distances below 4 Å or above 16 Å. The dihedral angle *ω* is formed by atoms C_α*i*_, C_β*i*_, C_β*j*_, and C_α*j*_, and the dihedral angle θ_*ij*_ is formed by atoms N_*i*_, C_α*i*_, C_β*i*_, and C_β*j*_. Both dihedral angles were discretized into 26 equal-sized bins in the range of [-180, 180°]. The planar angle *φ*_*ij*_ is formed by atoms C_α*i*_, C_β*i*_, and C_β*j*_. Planar angles were discretized into 26 equal-sized bins in the range of [0, 180°]. Orientation angles were not calculated for glycine residues, due to the absence of the C_β_ atom.

**Figure 1.**
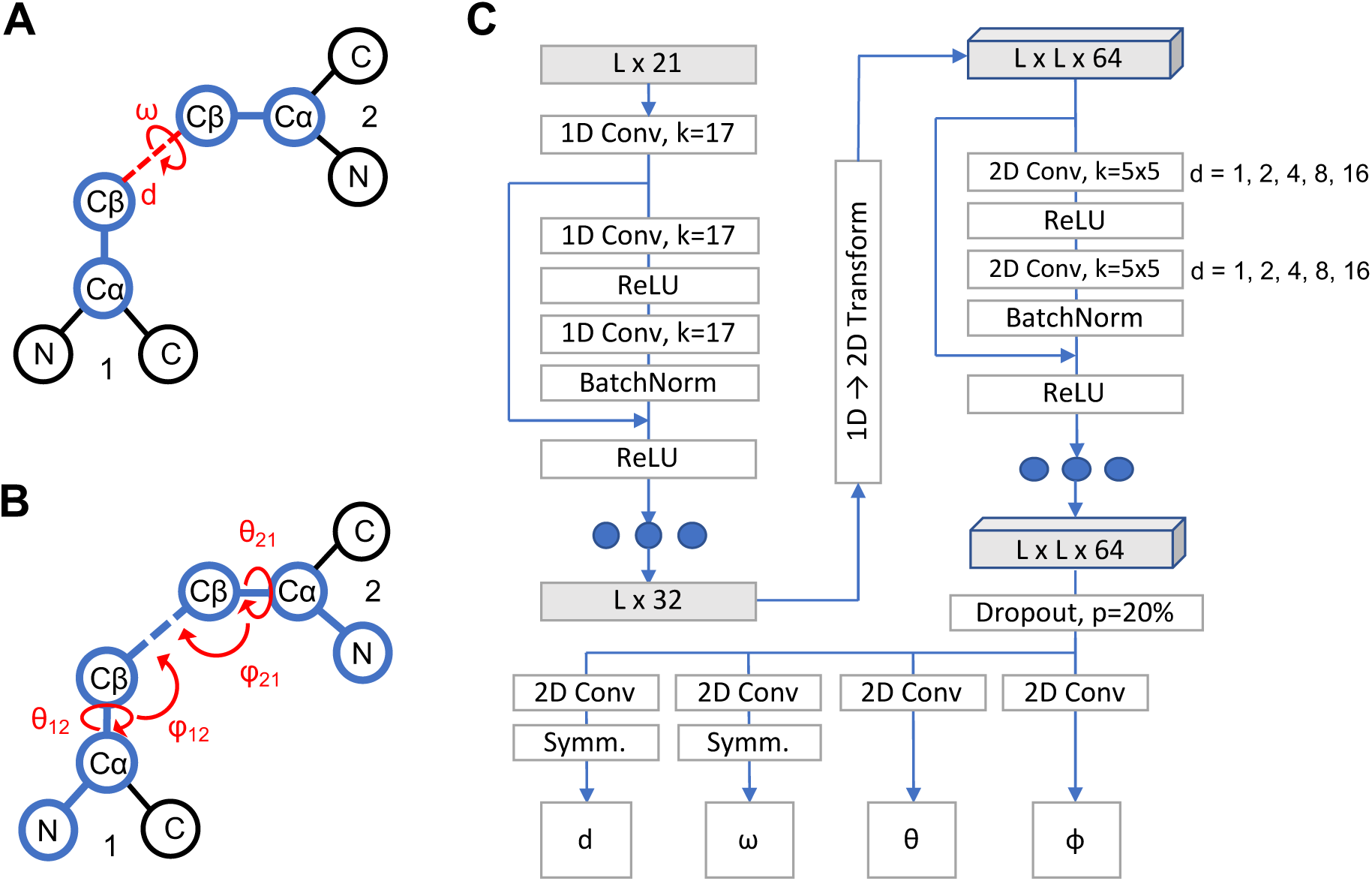
Architecture of DeepH3 deep residual neural network. **(A)** Illustration of the distance *d* and dihedral ω for two residues. **(B)** Illustration of the dihedrals θ_12_ and θ_21_ and planar angles ϕ_12_ and ϕ_21_ for two residues. **(C)** Architecture diagram of residual neural network to learn inter-residue geometries from concatenated antibody F_v_ chain sequences.

#### Network Architecture

DeepH3 applies a series of 1D and 2D convolutions to the aforementioned sequence input feature to predict four inter-residue geometries, as diagrammed in Figure 1C. The first 1D convolution (kernel size of 17) projects the *L*x21 input features up to an Lx32 tensor. Next, the Lx32 tensor passes through a set of three 1D residual blocks (two 1D convolutions with kernel size of 17), which maintain dimensionality. Following the 1D residual blocks, the sequential channels are transformed to pairwise by redundantly expanding the Lx32 tensor to dimension LxLx32 and concatenating with the transpose, resulting in a LxLx64 tensor. This tensor passes through 25 2D residual blocks (two 2D convolutions with kernel size of 5×5) that maintain dimensionality. Dilation of the 2D convolutions cycles through values of 1, 2, 4, 8, and 16 every five blocks (five cycles in total). Next, the network branches into four paths, which each apply a 2D convolution (kernel size of 5×5) to project down to dimension LxLx26 (for 26 output bins). Symmetry is enforced for the *d* and *ω* branches after the final convolution by summing the resulting tensor with its transpose. The four resulting LxLx26 tensors are converted to pairwise probability distributions for each output using the softmax function. DeepH3 was implemented using PyTorch [19] and is freely available at https://github.com/Graylab/deepH3-distances-orientations.

#### Training

Categorical cross-entropy loss was calculated for each output tensor and the resulting losses were summed with equal weight before backpropagation. The Adam optimizer was used with an initial learning rate of 0.01 and reduction of learning rate upon plateauing of total loss. Dropout was used after the last 2D residual block, with entire channels being zeroed out at 20% probability. The network was trained using 95% of antibody dataset described above (1,388 structures) for 30 epochs. Each epoch utilized the entire training dataset, with a batch size of 4. Training lasted about 35 hours using one NVIDIA Tesla K80 GPU on the Maryland Advanced Research Computing Center (MARCC).

### Network Predictions as Geometric Potentials

#### Implementation

To test the effectiveness of predicted geometric potentials in CDR H3 loop structure prediction, we collected Marze *et al.*’s RosettaAntibody set of 2,800 decoy structures for each of 49 antibody targets [14]. Marze *et al.* generated these structures by homology modeling, with decoys for each target assuming various heavy/light-chain orientations and non-H3 CDR loop conformations. Subsequently, we applied DeepH3 to each sequence in the benchmark dataset to produce pairwise probability distributions for the four output geometries. Distributions for pairs of residues that did not include a member of the CDR H3 loop were discarded. Additionally, pairs of residues for which the maximum probability bin of the distance output was greater than 12 Å were discarded to focus on local interactions that are likely to carry biophysical meaning. We also disregarded those predicted distributions that were not informative enough, chosen as those with a maximum probability below 10%. The remaining distributions were converted to potentials by taking the negative natural log of each output bin probability. Continuous, differentiable Rosetta constraints (`AtomPair` for *d*, `Dihedral` for *ω* and θ, and `Angle` for *φ*) were created for each potential using the built-in spline function. These constraint functions were applied to the decoy structures produced by RosettaAntibody to calculate a new DeepH3 energy term for each structure.

#### Discrimination Score

The discrimination score is a common metric for measuring the success of structure prediction calculations by assessing whether the minimum energy structures are near-native, with a lower value being indicative of a more successful prediction [15]. In order to compare between different energy schemes, we first scale the scores for all decoy structures such that the 95^th^ percentile energy has a value of 0.0 and the 5^th^ percentile energy has a value of 1.0. The discrimination score is then calculated as [20]:

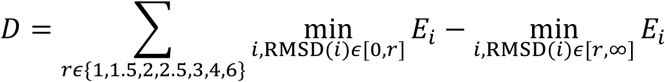

where *r* is the RMSD cutoff in Å, *E*_*i*_ is the scaled energy for the *i*-th decoy structure, and the discrimination score, *D*, is the sum of the energy differences for the best scoring decoys above and below each RMSD cutoff.

## Results

### DeepH3 Accurately Predicts Inter-Residue Geometries

To evaluate the accuracy of DeepH3’s predictions, we applied our model to the entire Rosetta antibody benchmark dataset (not seen during training or validation). For residue pairs involving a CDR H3 loop residue, the predicted values for each geometry are plotted against experimental structure values in Figure 2. DeepH3 displays effective learning across all outputs; the Pearson correlation coefficients (r) for d and ϕ were 0.87 and 0.79, respectively, and the circular correlation coefficients (r_c_) for dihedrals ω and θ were 0.52 and 0.88, respectively.

**Figure 2.**
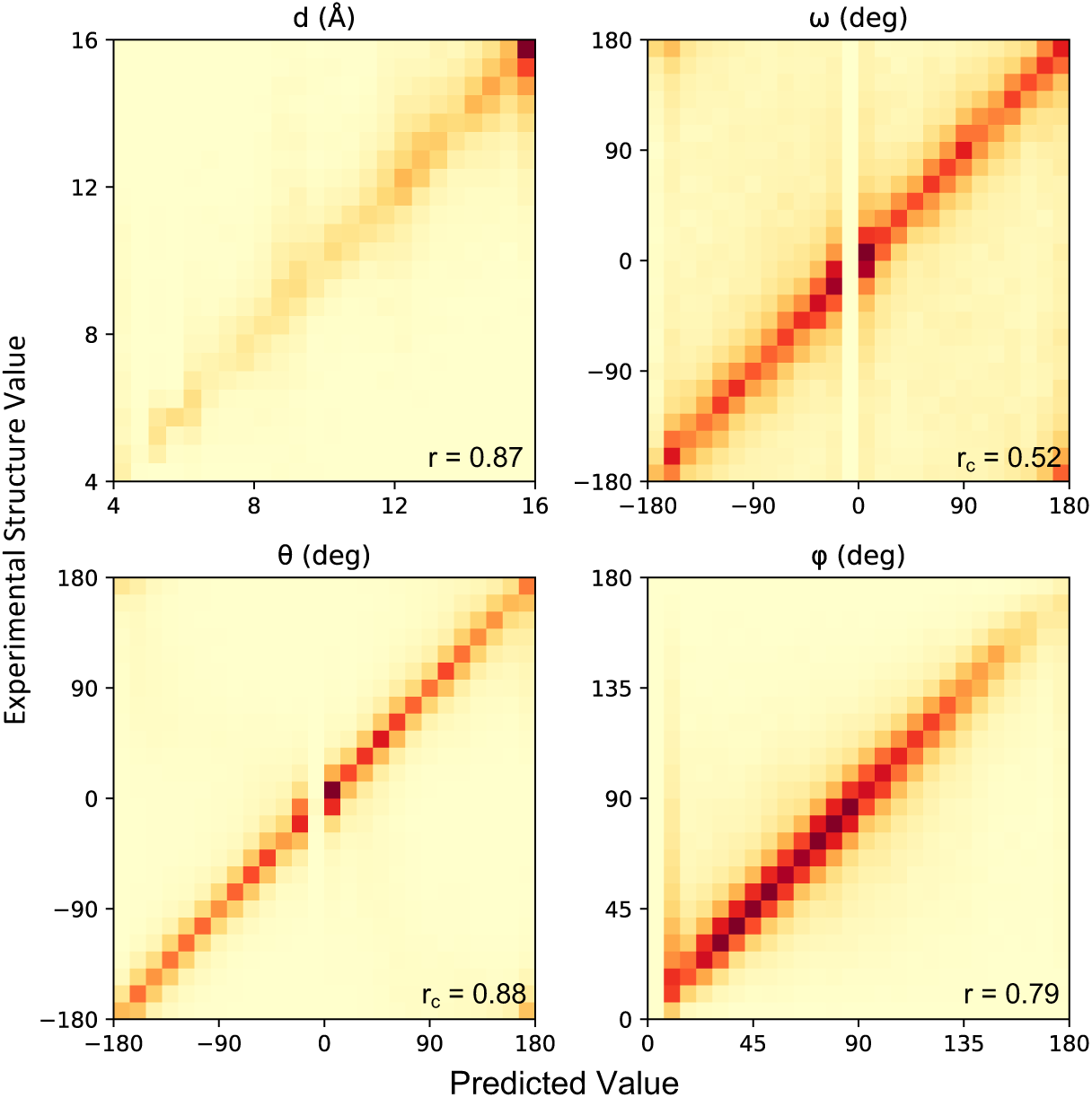
Accuracy of predicted inter-residue geometries. Pearson correlation coefficients (for d and ϕ) and circular correlation coefficients (for ω and θ) are calculated between DeepH3 predictions and experimental values.

### Geometric Potentials Effectively Discriminate Between Loop Structures

DeepH3 predictions were converted to geometric potentials (see Methods) and applied to RosettaAntibody generated structure decoys to evaluate the effectiveness of DeepH3 energy for identifying near-native structures. When the best-scoring structures (top 1) by Rosetta energy and DeepH3 energy were compared, DeepH3 selected better-, same-, and worse-RMSD structures for 30, 13, and 6 out of 49 targets, respectively, with an average RMSD improvement of 0.48 Å (Figure 3A). When the set of five best-scoring structures (top 5) by Rosetta energy and DeepH3 energy were considered, DeepH3 energy identified a better-, same-, and worse - RMSD structures for 26, 17, and 6 out of 49 targets, respectively, with an average RMSD improvement of 0.40 Å (Figure 3B). We also compared the ability of Rosetta energy and DeepH3 energy to discriminate between decoys for each benchmark target (Figure 3C). The mean discrimination scores for Rosetta energy and DeepH3 energy across the benchmark were −2.51 and −21.10, respectively, indicating that both methods were successful in general. When individual targets are considered, DeepH3 energy was successful in discriminating between decoys for 43 out of 49 targets, while Rosetta energy was successful for only 26 out of 49 targets.

**Figure 3.**
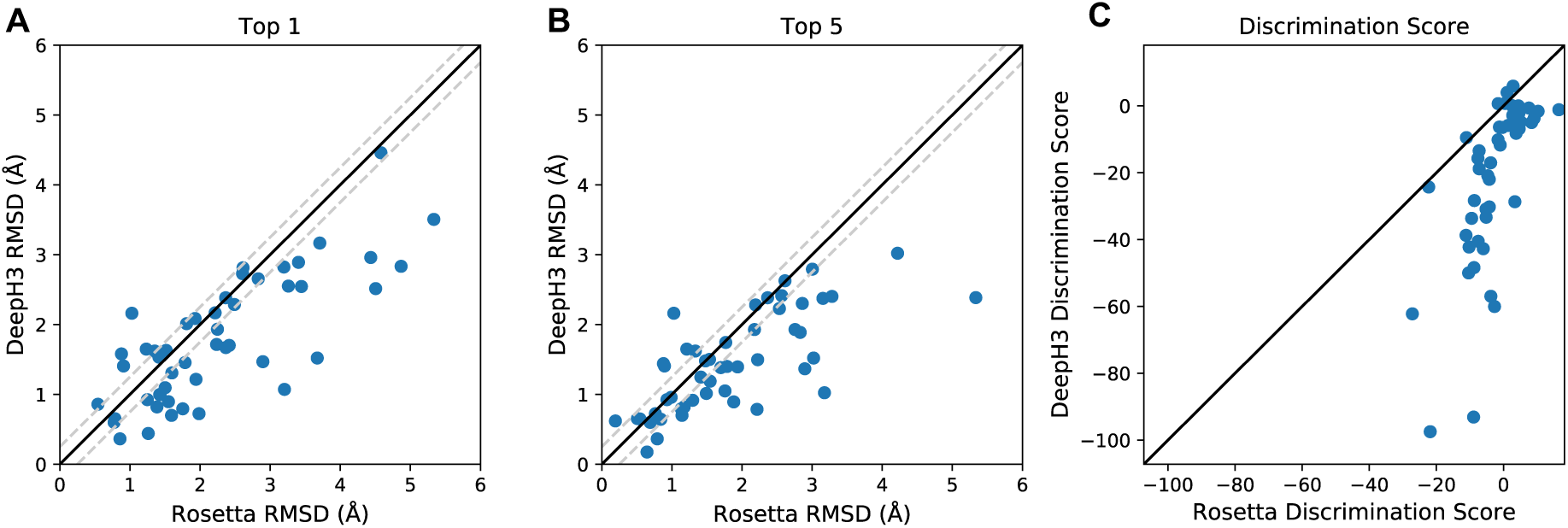
Effectiveness of predicted inter-residue geometries. **(A-B)** Comparison of the quality of structures selected by Rosetta energy and DeepH3 energy (using all geometric potentials). The quality of structures is considered the same if the difference in RMSD is within ±0.25 Å, indicated with dashed lines. **(A)** DeepH3 energy selected better-, same-, and worse-RMSD structures for 30, 13, and 6 out of 49 targets, respectively, when the best-scoring structures were compared (top 1). **(B)** When the set of five best-scoring structures were considered (top 5), DeepH3 energy identified better-, same-, and worse-RMSD structures for 26, 17, and 6 out of 49 targets, respectively. **(C)** Comparison of the discrimination scores for Rosetta energy and DeepH3 energy.

To provide a better understanding of how predicted geometric potentials improve discrimination between CDR H3 structures, we provide two case studies: anti-cytochrome C oxidase (mouse antibody with an eleven-residue CDR H3 loop, PDB ID: 1MQK) and sonepcizumab (humanized mouse antibody with a fourteen-residue CDR H3 loop, PDB ID: 3I9G) [15]. Figures 4A and 4C show energy funnels for anti-cytochrome C oxidase and sonepcizumab, respectively, with the discrimination score calculated for each. For anti-cytochrome C oxidase, Rosetta energy displays little ability to discriminate with structures ranging from 1 to 4 Å RMSD (D = 3.39). DeepH3 energy, however, earns a highly negative discrimination score (D = −28.68), indicating an ability to easily distinguish the near-native structures. The best-scoring 1mqk decoy structures as selected by Rosetta energy (orange, 3.20 Å RMSD) and DeepH3 energy (violet, 1.07 Å RMSD) are shown in Figure 4B. For sonepcizumab, Rosetta energy is generally successful in discriminating between decoys (D = −1.59), but again with minor energetic differences across a wide range of RMSD values. DeepH3 energy appears to converge to an alternative loop conformation around 2.5 Å RMSD, resulting in a poor discrimination score (D = 0.66). Figure 4D shows the best-scoring sonepcizumab decoy structures as selected by Rosetta energy (orange, 1.03 Å RMSD) and DeepH3 energy (violet, 2.16 Å RMSD).

**Figure 4.**
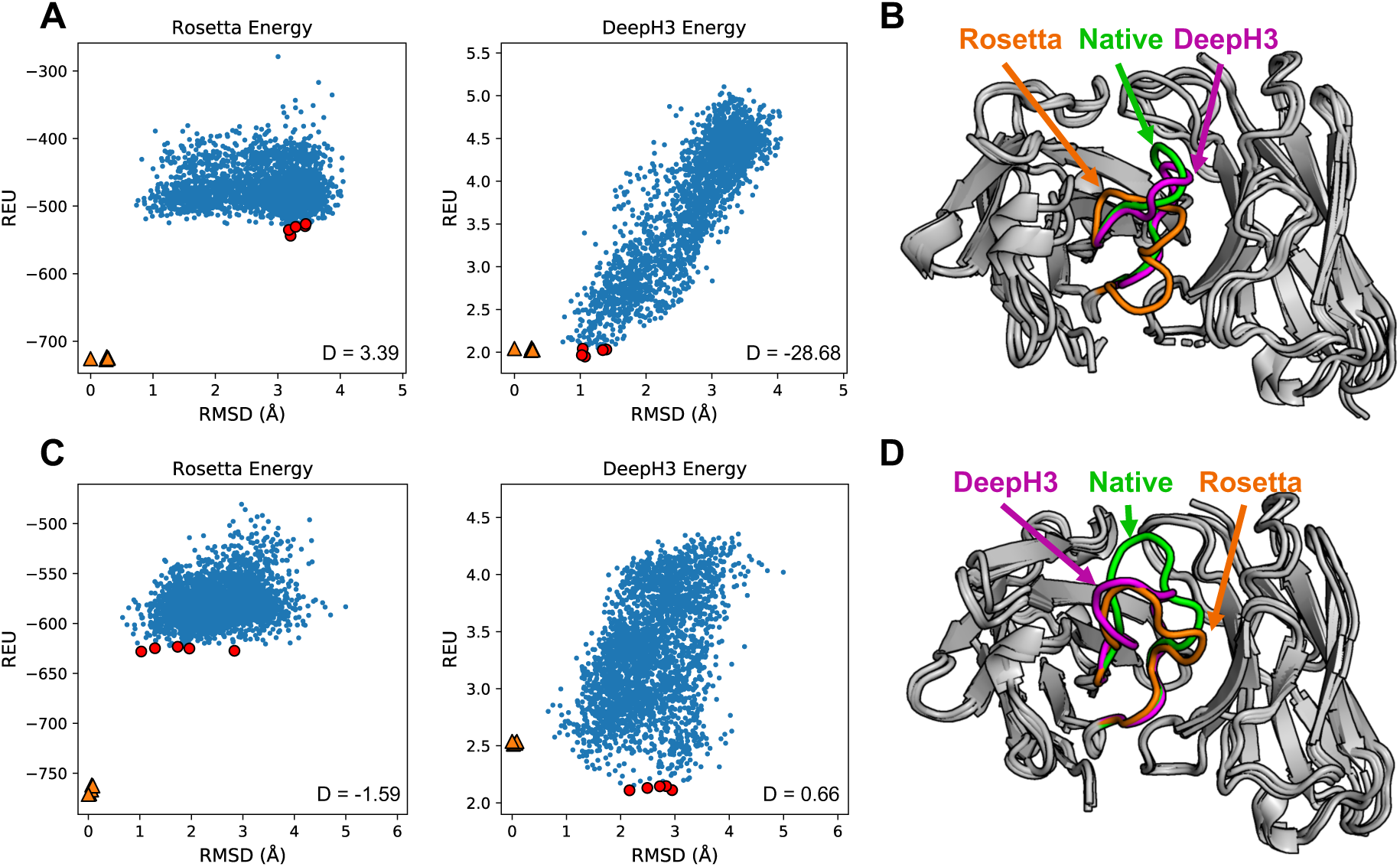
Results for two Rosetta antibody benchmark targets. **(A)** Plots of Rosetta energy and DeepH3 energy vs distance from the experimental structure for 2,800 decoy structures for anti-cytochrome C oxidase. The five best-scoring structures in each funnel plot are indicated in red. Five relaxed native structures are plotted as orange triangles. **(B)** Experimental structure of anti-cytochrome C oxidase (green) with best-scoring structures by Rosetta energy (orange, 3.20 Å RMSD) and DeepH3 energy (violet, 1.07 Å RMSD). **(C)** Plots of energy vs distance from the experimental structure for sonepcizumab. **(D)** Experimental structure of sonepcizumab (green) with best-scoring structures by Rosetta energy (orange, 1.03 Å RMSD) and DeepH3 energy (violet, 2.16 Å RMSD).

### Orientation Potentials are More Effective than Distance Potentials

We also evaluated the utility of individual geometric potentials for selecting low-RMSD decoys (Table 1) Notably, when DeepH3 distance potentials alone were used, performance was only moderately better than Rosetta energy. When the best-scoring structures by Rosetta energy and distance potentials were compared, distance potentials selected better-, same-, and worse-RMSD structures for 25, 13, and 11 out of 49 targets, respectively, with an average RMSD improvement of 0.34 Å. When the set of five best-scoring structures by Rosetta energy and distance potentials were considered, DeepH3 energy identified a better-, same-, and worse - RMSD structures for 20, 19, and 10 out of 49 targets, respectively, with an average RMSD improvement of 0.23 Å. Individual orientation potentials were more effective at selecting low-RMSD decoys than distance, even matching or outperforming the total DeepH3 energy by some metrics. We also calculated discrimination scores for each geometric potential (Table 2). Distance potentials display the weakest performance among geometric potentials but still show significant improvement over Rosetta energy, with 40 out of 49 simulations being successful and a mean discrimination score of −14.08. All three orientation potentials produced more successful simulations and lower mean discrimination scores than distance potentials.

**Table 1.** Performance of geometric potentials vs Rosetta energy function for selecting low-RMSD antibody decoys. Top-1 metrics compare the RMSD of the best-scoring structure by Rosetta energy against that of a given combination of DeepH3 potentials. Top-5 metrics compare the lowest-RMSD structure among the five best-scoring structures selected by Rosetta energy and that of a given DeepH3 potential. The average difference in RMSD between the structures selected by a given DeepH3 potential and Rosetta energy is reported as ΔRMSD (Å).

| Energy Terms | Top 1 |  |  |  | Top 5 |  |  |  |
| --- | --- | --- | --- | --- | --- | --- | --- | --- |
| | Better | Same | Worse | $\Delta\text{RMSD}$ | Better | Same | Worse | $\Delta\text{RMSD}$ |
| d | 25 | 13 | 11 | -0.34 | 20 | 19 | 10 | -0.23 |
| $\omega$ | 30 | 11 | 8 | -0.43 | 28 | 14 | 7 | -0.39 |
| $\theta$ | 31 | 11 | 7 | -0.50 | 26 | 17 | 6 | -0.39 |
| $\phi$ | 29 | 14 | 6 | -0.46 | 26 | 17 | 6 | -0.41 |
| d + $\omega$ + $\theta$ + $\phi$ | 30 | 13 | 6 | -0.48 | 26 | 17 | 6 | -0.40 |

**Table 2.** Discrimination score metrics for Rosetta energy and DeepH3 potentials. Negative discrimination scores, *D*, are considered successful and positive are considered unsuccessful.

| Energy Terms | Successful | Unsuccessful | Mean $D$ |
| --- | --- | --- | --- |
| Rosetta Energy | 29 | 20 | -2.5 |
| d | 40 | 9 | -14.1 |
| $\omega$ | 44 | 5 | -15.1 |
| $\theta$ | 43 | 6 | -28.1 |
| $\phi$ | 45 | 4 | -17.9 |
| d + $\omega$ + $\theta$ + $\phi$ | 43 | 6 | -21.1 |

## Discussion

The results here suggest that the significant advances by deep learning approaches in general protein structure can also be realized in subproblems in structural modeling. Specifically, we demonstrate that a deep residual network can effectively capture the local inter-residue interactions that define antibody CDR H3 loop structure. DeepH3 achieves these results without co-evolutionary data, while using significantly fewer residual blocks (3 1D + 25 2D blocks) than similar networks, such as AlphaFold (220 2D blocks) [10], RaptorX (6 1D + 60 2D blocks), and trRosetta (61 2D blocks). Fewer blocks may suffice because we limited our focus to antibodies, which are highly conserved, rather than the entire universe of protein structures. In the future, similar specialized networks could achieve enhanced performance in other challenging domains of protein structure prediction.

Breakdown of DeepH3 energy into individual geometric potentials revealed that inter-residue orientations were significantly more effective for scoring CDR H3 loop structure than distances. This finding was surprising, given the improvements that distances alone have enabled in general protein structure prediction. It is possible that while accurate distance predictions are effective at placing residues globally, the attention to local interactions necessary for CDR H3 loop modeling is better captured by inter-residue orientations. While this work only utilized predicted inter-residue geometries to discriminate between CDR H3 loop structures, we expect that a similar approach would yield improvement in the generation of structures as well.

## Acknowledgments

We thank Jeliazko Jeliazkov for helpful discussions and his RosettaAntibody expertise. Computational power was provided by the Maryland Advanced Research Computing Center (MARCC).

## References

[1] C. Chothia, A. M. Lesk, A. Tramontano, M. Levitt, S. J. Smith-Gill, G. Air and S. Sheriff, “Conformations of immunoglobulin hypervariable regions.,” Nature, vol. 342, no. 6252, p. 877, 1989.

[2] B. North, A. Lehmann and R. L. Dunbrack, “A new clustering of antibody CDR loop conformations.,” Journal of molecular biology, vol. 406, no. 2, pp. 228–256, 2011.

[3] B. D. Weitzner, D. Kuroda, N. Marze, J. Xu and J. J. Gray, “Blind prediction performance of RosettaAntibody 3.0: grafting, relaxation, kinematic loop modeling, and full CDR optimization.,” Proteins: Structure, Function, and Bioinformatics, vol. 82, no. 8, pp. 1611–1623, 2014.

[4] J. C. Almagro, A. Teplyakov, J. Luo, R. W. Sweet, S. Kodangattil, F. Hernandez-Guzman and G. L. Gilliland, “Second antibody modeling assessment (AMA-II).,” Proteins: Structure, Function, and Bioinformatics, vol. 82, no. 8, pp. 1553–1562, 2014.

[5] M. Fasnacht, K. Butenhof, A. Goupil-Lamy, F. Hernandez-Guzman, H. Huang and L. Yan, “Automated antibody structure prediction using Accelrys tools: Results and best practices.,” Proteins: Structure, Function, and Bioinformatics, vol. 82, no. 8, pp. 1583–1598, 2014.

[6] J. K. Maier and P. Labute, “Assessment of fully automated antibody homology modeling protocols in molecular operating environment.,” Proteins: Structure, Function, and Bioinformatics, vol. 82, no. 8, pp. 1599–1610, 2014.

[7] H. Shirai, K. Ikeda, K. Yamashita, Y. Tsuchiya, J. Sarmiento, S. Liang, T. Morokata, K. Mizuguchi, J. Higo, D. M. Standley and H. Nakamura, “High-resolution modeling of antibody structures by a combination of bioinformatics, expert knowledge, and molecular simulations.,” Proteins: Structure, Function, and Bioinformatics, vol. 82, no. 8, pp. 1624–1635, 2014.

[8] K. Zhu, T. Day, D. Warshaviak, C. Murrett, R. Friesner and D. Pearlman, “Antibody structure determination using a combination of homology modeling, energy-based refinement, and loop prediction.,” Proteins: Structure, Function, and Bioinformatics, vol. 82, no. 8, pp. 1646–1655, 2014.

[9] M. Berrondo, S. Kaufmann and M. Berrondo, “Automated Aufbau of antibody structures from given sequences using Macromoltek’s SmrtMolAntibody.,” Proteins: Structure, Function, and Bioinformatics, vol. 82, no. 8, pp. 1636–1645, 2014.

[10] A. Senior, R. Evans, J. Jumper, J. Kirkpatrick, L. Sifre, T. Green, C. Qin and H. Penedones, “Improved protein structure prediction using potentials from deep learning.,” Nature, 2020.

[11] J. Xu, “Distance-based protein folding powered by deep learning.,” Proceedings of the National Academy of Sciences, vol. 116, no. 34, pp. 16856–16865, 2019.

[12] J. Yang, I. Anishchenko, H. Park, Z. Peng, S. Ovchinnikov and D. Baker, “Improved protein structure prediction using predicted interresidue orientations.,” Proceedings of the National Academy of Sciences, 2020.

[13] S. Wang, S. Sun, Z. Li, R. Zhang and J. Xu, “Accurate de novo prediction of protein contact map by ultra-deep learning model.,” PLoS computational biology, vol. 13, no. 1, p. e1005324, 2017.

[14] N. A. Marze, S. Lyskov and J. J. Gray, “Improved prediction of antibody VL–VH orientation.,” Protein Engineering, Design and Selection, vol. 29, no. 10, pp. 409–418, 2016.

[15] B. D. Weitzner and J. J. Gray, “Accurate structure prediction of CDR H3 loops enabled by a novel structure-based C-terminal constraint.,” The Journal of Immunology, vol. 198, no. 1, pp. 505–515, 2017.

[16] J. Adolf-Bryfogle, Q. Xu, B. North, A. Lehmann and R. L. Dunbrack Jr., “PyIgClassify: a database of antibody CDR structural classifications.,” Nucleic acids research, vol. 43, no. D1, pp. D432–D438, 2014.

[17] J. Dunbar, K. Krawczyk, J. Leem, T. Baker, A. Fuchs, G. Georges, J. Shi and C. M. Deane, “SAbDab: the structural antibody database.,” Nucleic acids research, vol. 42, no. D1, pp. D1140–D1146, 2013.

[18] S. Wang, S. Sun and J. Xu, “Analysis of deep learning methods for blind protein contact prediction in CASP12.,” Proteins: Structure, Function, and Bioinformatics, vol. 86, pp. 67–77, 2018.

[19] A. Paszke, S. Gross, F. Massa, A. Lerer, J. Bradbury, G. Chanan, T. Killeen, Z. Lin, N. Gimelshein, L. Antiga and A. Desmaison, “PyTorch: An imperative style, high-performance deep learning library.,” in Advances in Neural Information Processing Systems, 2019.

[20] P. Conway, M. D. Tyka, F. DiMaio, D. E. Konerding and D. Baker, “Relaxation of backbone bond geometry improves protein energy landscape modeling,” Protein Science, vol. 21, no. 3, pp. 47–55, 2014.

